# Musicianship modulates cortical (but not brainstem) effects of attention on processing musical triads

**DOI:** 10.1101/2024.09.21.614251

**Authors:** Jessica MacLean, Elizabeth Drobny, Rose Rizzi, Gavin M. Bidelman

## Abstract

**Background:** Many studies have demonstrated benefits of long-term music training (i.e., musicianship) on the neural processing of sound, including simple tones and speech. However, the effects of musicianship on the encoding of simultaneously presented pitches, in the form of complex musical chords, is less well-established. Presumably, musicians’ stronger familiarity and active experience with tonal music might enhance harmonic pitch representations, perhaps in an attention-dependent manner. Additionally, attention might influence chordal encoding differently across the auditory system. To this end, we explored the effects of long-term music training and attention on processing of musical chords at brainstem and cortical levels.

**Method:** Young adult participants were separated into musician and nonmusician groups based on extent of formal music training. While recording EEG, listeners heard isolated musical triads that differed only in the chordal third: major, minor, and detuned (4% sharper third from major). Participants were asked to correctly identify chords via key press during active stimulus blocks and watched a silent movie during passive blocks. We logged behavioral identification accuracy and reaction times and calculated information transfer based on the behavioral chord confusion patterns. EEG data were analyzed separately to distinguish between cortical (event-related potential, ERP) and subcortical (frequency-following response, FFR) evoked responses.

**Results:** We found musicians were (expectedly) more accurate, though not faster, than nonmusicians in chordal identification. For subcortical FFRs, responses showed stimulus chord effects but no group differences. However, for cortical ERPs, whereas musicians displayed P2 (∼150 ms) responses that were invariant to attention, nonmusicians displayed earlier and reduced P2 during passive listening. Listeners’ degree of behavioral information transfer (i.e., success in distinguishing chords) was also better in musicians and correlated with their neural differentiation of chords, assessed via pairwise differences in the ERPs.

**Conclusions:** Our data suggest long-term music training strengthens even the passive cortical processing of musical sounds, supporting more automated brain processing of musical chords with less reliance on attention. Our results also suggest the degree to which listeners can behaviorally distinguish chordal triads is directly related to their neural specificity to musical sounds primarily at cortical rather than subcortical levels.

## INTRODUCTION

Many studies have demonstrated benefits of long-term music training (i.e., musicianship) for neural processing of sound, including simple tones [1] and speech [2]. This “musician enhancement” in brain processing has been linked to a variety of auditory perceptual skills, especially those related to spectral processing. For example, musicians show improved abilities to detect violations in musical pitch patterns [3-5], smaller frequency difference limens [6-8], and reduced susceptibility to timbral influences on pitch perception [9]. Musicians’ auditory benefits also confer improvements to non-musical sounds, including speech [10-14]. Such enhancements to behavioral and neural indicators of sound processing are important for understanding how music engagement may support everyday listening skills.

Despite a wealth of cross-domain investigations into the influence of music experience on neural speech encoding, less is known about the effects of music training on neural encoding of *musical* sounds. Given that music performance necessitates the precise manipulation and monitoring of pitch information (e.g., to remain in tune within an ensemble), musicianship may engender superior musical pitch encoding abilities. Bidelman et al. [1] investigated neural processing of arpeggiated triad chords, in which one note was played at a time (i.e., melodically). Major, minor, and detuned (±4% frequency) triads only differed in the chordal third, the middle note of which determined the perceived quality of the chord. Major and minor chords are often heard in Western music and can be reliably distinguished by individuals without music training as sounding “happy” and “sad,” respectively [15]. In contrast, the detuned chord represented a triad less familiar to participants given its absence from common Western scales. Bidelman et al. [1] found musicians’ frequency-following responses (FFRs), a neurophonic potential reflecting subcortical (brainstem) phase locking, showed stronger neural onset synchronization and encoding of the chordal third than nonmusicians’, regardless of whether the chord was in or out of tune. Musicians’ enhanced FFRs were paralleled by their improved performance on pitch discrimination tasks. While musicians showed superior discrimination of three chord variants, nonmusicians were only sensitive to major/minor distinctions and could not detect more subtle chord mistuning. Though this study suggests musicianship strengthens passively-evoked brainstem responses to musical triads, it does not elucidate whether such experience-dependent enhancements are uniform at different levels of auditory processing, nor how they change under different states of attention. Here, we investigated the effects of musicianship on the neural processing of harmonic musical chords at multiple levels of the auditory system.

In this vein, several studies have shown musician enhancements to speech and music at a cortical level utilizing event-related potentials (ERPs) [16-20]. However, cortical neuroplasticity associated with music training is sometimes observed in the absence of subcortical enhancements [21]. This raises the question of where in the auditory system music training exerts its neuroplastic influences on auditory coding and which level of processing is most predictive of musicians’ behavioral benefits for musical pitch. Relatedly, the effects of musicianship may be attention-dependent. Attention can modulate both subcortical FFRs [22,23] and cortical ERPs to speech [24,25]. However, most studies demonstrating FFR enhancements in musicians have utilized only passive listening paradigms where participants watch a silent movie and do not interact with the stimuli [2]. At least for speech sounds, musicians’ FFR enhancements are not necessarily observed under active listening even when auditory cortical potentials do show attentional gains [21]. Thus, although musicianship may exert influences on attention at the behavioral level [10,26], it may do so with differing degrees across the auditory brain hierarchy.

Given the equivocal nature of the literature, the current study aimed to test whether brainstem FFRs and cortical ERPs are sensitive to attention and reflect a neural correlate of active music perception. We used musical chord triads [cf. 1] in an active identification paradigm [23,27,28] designed to simultaneously record FFRs/ERPs with behavioral responses. Chordal stimuli included the major and minor triads of Western music and a detuned (4% sharp) version. Chords were advantageous to our design because their perceptual identification required listeners to correctly “hear out” only a single middle note (i.e., chordal third) flanked by two other pitches. We hypothesized the chordal third would be selectively enhanced in neural responses when listeners correctly identified the chord. We also measured listeners with a range of musical training as we hypothesized musically experienced listeners would be more successful at musical chord identification tasks [e.g., 1] and thus show stronger attentional changes in their FFRs and/or ERPs.

## MATERIALS AND METHODS

### Participants

Our sample included N = 15 young adults (age range= 19-31 years, 10 female/5 male). All participants were monolingual English-speakers and had normal hearing (pure tone thresholds ≤25 dB HL; octave frequencies 250-8000 Hz). Participants had an average of 17.6 ± 2.80 years of education. Based on participants’ self-reported musical training (mean= 9.47 ± 8.26 years; range= 0-24 years), participants were split into two groups: musicians (M; n= 9) had greater than 5 years of training and nonmusicians (NM; n =6) had fewer than 5 years of training (see [29] for a similar definition of “musician”). Each participant provided written, informed consent in compliance with a protocol approved by the Institutional Review Board of Indiana University and was paid for their time.

### Stimuli and task

We used harmonic chord stimuli to evoke FFRs [e.g., 1]. Each chord consisted of three concurrent pitches (root, 3^rd^, 5^th^) that differed only in the chordal third (i.e., middle pitch). Consequently, category labeling could only be accomplished if listeners were able to perceptually “hear out” the chord-defining pitch. Two chords were exemplary of Western music practice (major and minor chord); the third represented a detuned version in which the chord’s third was made sharp (4%) from its major version. The minor and major third intervals connote the valence of “sadness” and “happiness” even to non-musicians [30] and are easily described to listeners unfamiliar with music-theoretic labels [31]. A 4% deviation is greater than the just-noticeable difference for frequency (<1%) [32] but smaller than a full semitone (6%). This amount of deviation is similar to previously published reports examining musicians’ and non-musicians’ EEG responses to detuned triads [e.g., 1,4,30]. Individual chord notes were synthesized using complex-tones (6 iso-amplitude harmonics added in sine phase; 100 ms duration; 5 ms ramp). The fundamental frequency (F0) of each of the three notes (i.e., root, 3^rd^, 5^th^) per triad were as follows: major = 220, <u>277</u>, 330 Hz; minor = 220, <u>262</u>, 330 Hz; detuned = 220, <u>287</u>, 330 Hz. Critically, the F0 (indeed all spectral cues) in our stimuli exceeded 220 Hz, which is substantially higher than the phase-locking limits of cortical neurons observed in any animal or human studies [33-35]. This ensured our FFRs were of brainstem origin [21,28,35]. Before initiating the task proper, participants were allowed to freely replay the three stimuli for familiarization.

To efficiently record FFRs during an online behavioral task while obtaining the high (i.e., several thousand) trial counts needed for response visualization, we used a clustered interstimulus interval (ISI) presentation paradigm [23,27,28]. Single tokens were presented in blocks of 15 repetitions with a rapid ISI (10 ms). After the clustered block of tokens ended, the ISI was slowed to 1500 ms and a single token was presented to cue the behavioral response and allow for recording of an un-adapted cortical ERP [27]. Participants then indicated their percept (major, minor, detuned) as quickly and accurately as possible via the keyboard. Following the behavioral response and a period of silence (250 ms), the next trial cluster commenced. Per token, this paradigm allowed 1980 presentations for input to FFR analysis and 66 ERP presentations and behavioral responses. We calculated percent correct identification per stimulus condition.

In addition to the active condition, we ran a passive block that did not require an overt task. For passive presentation, participants were instructed to ignore the sounds they heard and watch a self-selected movie with subtitles to maintain a calm and wakeful state [19]. Because there was no behavioral response in passive blocks, we included an ISI of 600 ms between token clusters (i.e., comparable the estimated reaction time (RT) in the active block) to ensure that the overall pacing of stimulus delivery was comparable between listening conditions. The order of active and passive blocks was randomized.

Stimulus presentation was controlled via MATLAB (The MathWorks, Natick, MA, USA) routed to a TDT RZ6 (Tucker-Davis Technologies, Alachua, FL, USA) signal processor. Stimuli were presented binaurally at 80 dB SPL with rarefaction polarity. We used shielded insert headphones (ER-2; Etymotic Research) to prevent pickup of electromagnetic artifacts from contaminating neural recordings [36,37].

### EEG recording procedures

We used Curry 9 software (Compumedics Neuroscan, Charlotte,NC) and Neuroscan Synamps RT amplifiers to record the EEG data. Continuous EEGs were recorded differentially between scalp Ag/AgCl disk electrodes placed on the high forehead at the hairline (∼Fpz) referenced to linked mastoids (M1/M2); a mid-forehead electrode served as ground. This montage is optimal for pickup of the vertically oriented FFR dipole in the midbrain [38]. Electrode impedances remained ≤ 5 kΩ throughout the duration of recording. EEGs were digitized at 10 kHz to capture the fast activity of FFR. Trials exceeding a ±120 µV threshold were automatically rejected from the average. We separately band-pass filtered responses from 200-2500 Hz and 1-30 Hz (zero-phase Butterworth filters; slope = 48 dB/octave) to isolate FFRs and ERPs, respectively [39,40]. Data were then epoched relative to the time-locking trigger for each response class (i.e., FFR: 0-105 ms, ERP: -200-1000 ms), pre-stimulus baselined, and averaged for each stimulus token. Data preprocessing was performed in BESA Research v7.1 (BESA, GmbH, Gräfelfing, Germany).

### Brainstem FFR analysis

From each FFR waveform, we generated Fast Fourier Transforms (FFTs) to quantify the spectral information in each response. Amplitudes of the spectral peak corresponding to the chordal 3^rd^ (227-287 Hz) were identified in all conditions by two independent observers. Inter-rater reliability across all measurements was exceptional [*r* = 0.99, *p*<0.0001]. F0—related to pitch encoding— represents the dominate energy in the FFR and can be modulated by attention and listeners’ trial-by-trial categorical hearing [22,23,37,41]. We focused on the F0 of the chordal 3^rd^ as this was the category-defining pitch of each stimulus. Onset latency and amplitude were measured as the first positive peak in the FFR waveform in the 7-15 ms time window, the expected onset latency of brainstem responses [42,43].

### Cortical ERP analysis

From ERP waveforms to each stimulus, we measured the amplitude and latency of the N1 and P2 deflections between 100-140 ms and 135-175 ms, respectively. The analysis window was guided by visual inspection of the grand-averaged data. We selected N1 due to its well-documented susceptibility to attention effects and musicianship [44-50] and P2 for its sensitivity to long-term plasticity from musicianship [19,21].

### Confusion matrices and Information Transfer (IT) analysis

To examine the pattern of behavioral responses to chordal stimuli (in the active task) we first computed confusion matrices [51]. We then used information transfer (IT) [52] to directly assess the correspondence between stimulus input and behavioral output. IT is defined as the ratio of transmitted information between *x* and *y* [i.e., *T*(*x;y*)] to the input entropy (*H*_*x*_), expressed as a percentage. *T*(*x;y*) represents the transmission of information (in an information theoretic sense) from *x* to *y*, measured in bits per stimulus, and was computed from confusion matrices via Eq. 1:

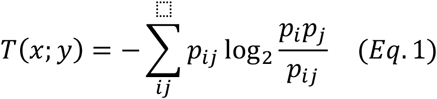

where *p*_*i*_, and *p*_*j*_, are the probabilities of the observed input and output variables, respectively, and *p*_*ij*_ is the joint probability of occurrence for observing input *i* with output *j*. These probabilities were computed from the confusion matrices as *p*_*i*_*= n*_*i*_/*N, p*_*j*_*= n*_*j*_/*N*, and *p*_*ij*_*= n*_*ij*_/*N*, where *n*_*i*_ is the frequency of stimulus *i, n*_*j*_ is the frequency of response *j*, and *n*_*ij*_ is the frequency of the joint occurrence of stimulus *i* and response *j* (i.e., diagonal elements of the confusion matrix) in the sample of *N* total observations (Miller & Nicely, 1955). The input entropy *H*_*x*_ is given by Eq. 2:

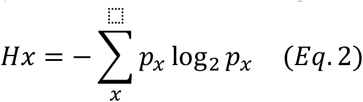

In the current study, all stimuli occurred with equal probability (i.e., *p*_*x*_ = 0.33). IT was then computed as (Eq. 3):

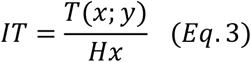

This metric varies from 0 to 100%. Intuitively, if the transmission is poor and a listener’s response does not closely correlate with the stimulus, then *IT* will approach zero; alternatively, if the response can be accurately predicted from the stimulus then *IT* will approach unity (i.e., 100% information transfer). *IT* was computed from the behavioral chordal confusion matrices for each participant and then group averaged. Comparisons of *IT* allowed us to assess the degree to which stimulus information was faithfully transmitted to perception in each listener.

### Neural differentiation of musical chords

We reasoned that listeners with more robust perceptual identification and fewer behavioral confusions would show maximally differentiable cortical ERPs across chordal stimuli [51]. To this end, we quantified the dissimilarity in listeners’ ERPs across the three chordal triads via the pairwise difference in ERP N1-P2 amplitude between chords (i.e., major-minor, major-detuned, minor-detuned). We focused on the N1-P2 for this analysis because this complex represents the overall magnitude of the auditory cortical ERPs and previous work has shown robust neural decoding of auditory stimulus properties in the time window of the N1-P2 deflections [51,53]. Differences were computed via the Euclidean distance between peak-to-peak amplitude values (MATLAB *pdist2* function). We then averaged the three distances (per participant) to quantify the overall differentiation of stimuli in their ERPs.

### Statistical analysis

Unless otherwise noted, we analyzed dependent variables using mixed model ANOVAs in the R (version 4.2.2) [54] lme4 package [55]. To analyze behavioral outcomes of accuracy (percent correct, PC) and information transfer [IT; 52], we modeled fixed effects of group (2 levels; M v. NM) and stimulus (3 levels; major v. minor v. detuned), with a random effect of subject. To analyze neural outcomes (FFR onset latency and amplitude, ERP P2 latency and amplitude), we added an additional fixed effect of condition (2 levels; active v. passive). Effect sizes are reported as partial eta squared 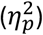 and degrees of freedom (*d*.*f*.) using Satterthwaite’s method. A priori significance level was set at α = 0.05. We examined brain-behavior correspondences via Pearson correlations between behavioral IT and ERP distances. Though statistical analyses are reported here with music as a fixed categorical variable, treating years of music training as a continuous predictor did not substantially change the results.

## RESULTS

### Behavioral chord identification

**Figure 1** displays behavioral results for all participants separated by musician (M) and nonmusician (NM) groups across all three chordal stimuli (major, minor, detuned). An ANOVA on behavioral accuracy revealed an interaction between group and stimulus [*F*(2, 26) = 3.61, *p* = 0.041, 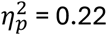]. Post hoc Tukey-adjusted pairwise comparisons revealed that whereas Ms performed equally well identifying all three chords (all pairwise comparisons: *p* > 0.77), NMs shows a gradient performance with better identification of the minor relative to other chords (**Fig. 1A**). These data suggest that musicians were more accurate (but not faster) in identifying musical chords. Behavioral confusion matrices (**Figure 1B**) for all stimuli were used to calculate information transfer (IT) [52] for both groups (**Figure 1C**). A *t*-test between groups indicated significantly higher IT for Ms relative to NMs [*t*(13) = 6.15, *p* < 0.001].

**Figure 1.**
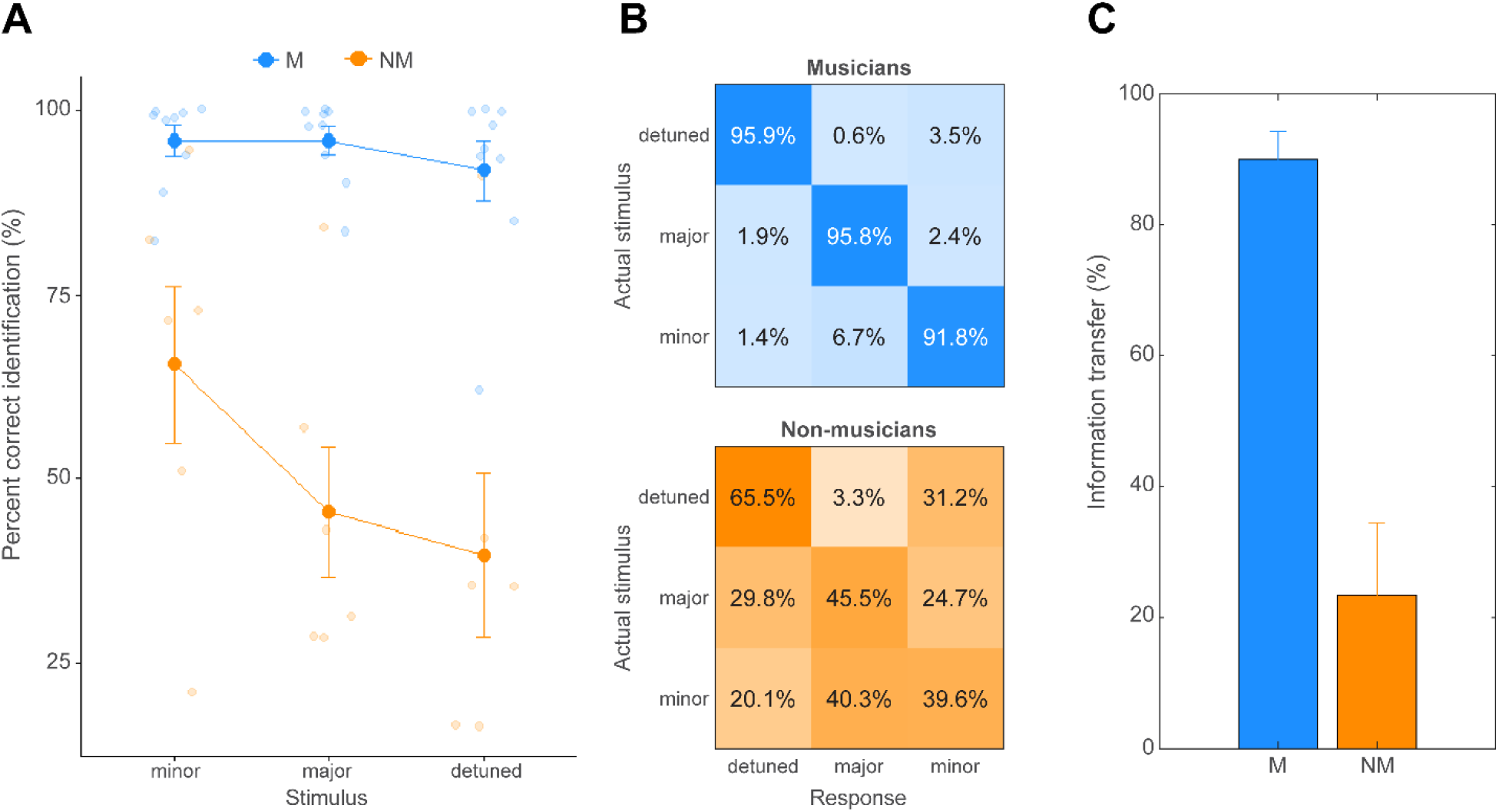
Behavioral performance in chord identification for musicians and nonmusicians (active condition). **(A)** Musicians outperformed nonmusicians in identification accuracy across the board. Nonmusicians showed more gradient performance and were better at identifying minor relative to the major and detuned chords. **(B)** Mean behavioral confusion matrices per group. Cells denote the proportion of responses labelled as a given chord relative to the actual stimulus category presented. Diagonals show correct responses. Chance level=33%. NMs show more perceptual confusions than Ms, especially between major and detuned chords. **(C)** Information transfer (IT) derived from the perceptual confusion matrices. IT represents the degree to which listeners’ responses can be accurately predicted given the known input stimulus [52]. Values approaching 100% indicate perfect prediction of the response given the input; values approaching 0 indicate total independence of the stimulus and response. IT is near ceiling and much larger in Ms indicating higher fidelity differentiation of musical stimuli. Error bars = ± 1 S.E.M.

### Brainstem FFRs

**Figure 2** shows subcortical FFRs to all three chordal stimuli for both groups and attention conditions. FFR response metrics are shown in **Figure 3**. An ANOVA on FFR onset amplitude revealed a significant main effect of stimulus [*F*(2, 65) = 7.55, *p* = 0.0011, 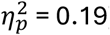] (**Fig. 3A**). Post hoc Tukey adjusted comparisons revealed that stimulus effect was driven by stronger onset amplitudes for the detuned relative to the minor (*p* = 0.0008) and major (*p* = 0.052) chords, respectively. There were no other main nor interaction effects (all *ps*> 0.3). FFR onset latency did not vary with attention condition [*F*(1, 65) = 0.71, *p* = 0.40, 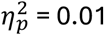], stimulus [*F*(2, 65) = 2.13, *p* = 0.13, 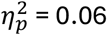], group [*F*(1, 13) = 0.14, *p* = 0.71, 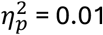], nor their two- and three-way interactions (all *ps* > 0.60; **Fig. 3B**).

**Figure 2.**
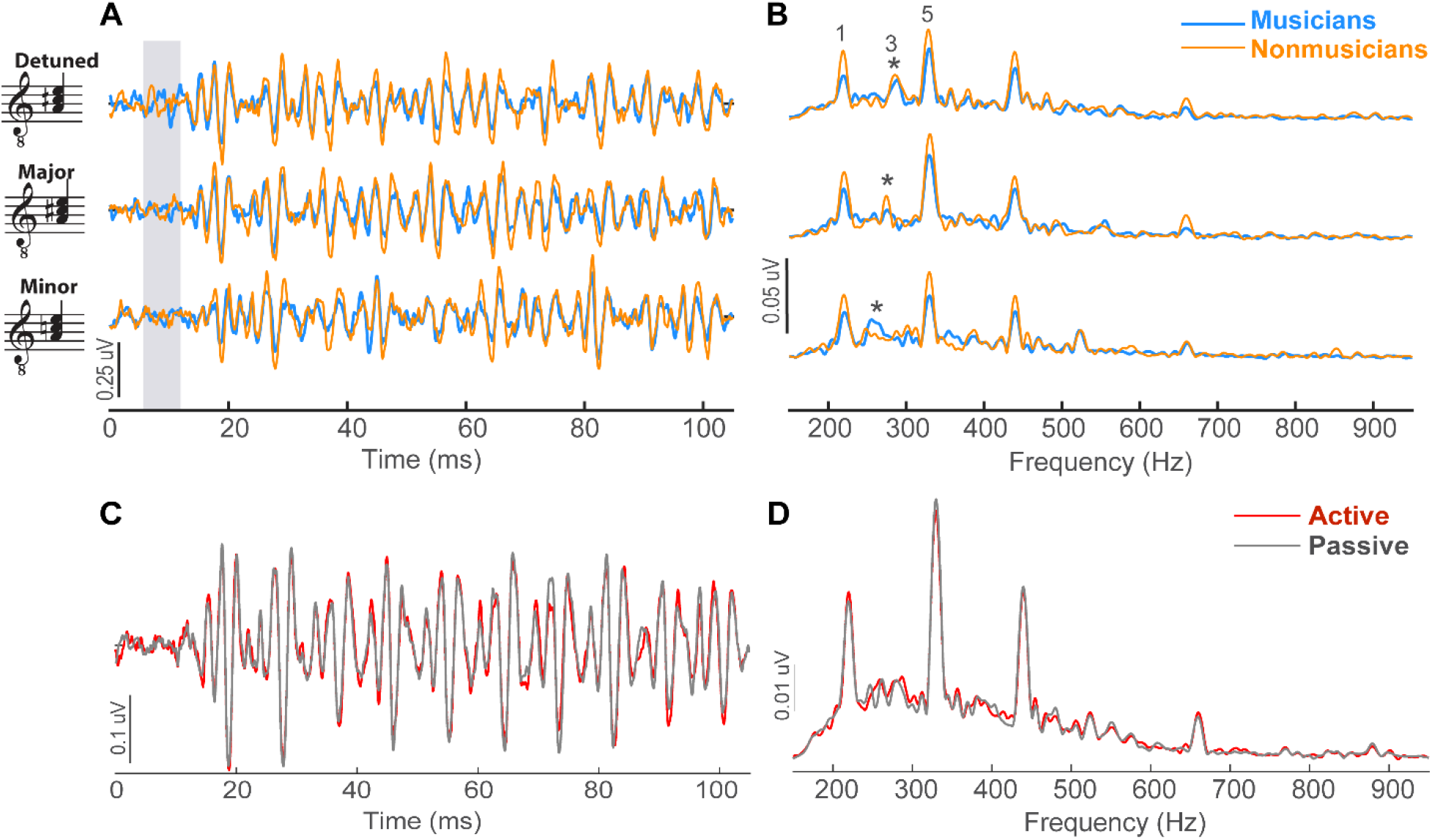
Group averaged FFRs to chordal stimuli. **(A)** Time waveforms to chordal stimuli in musicians and nonmusicians (active condition only). **(B)** Response spectra (FFTs). FFRs did not differ between groups. Peaks for the chordal root (1; 220 Hz), third (3*; 262-287 Hz) and fifth (5; 330 Hz) are demarcated. Only the chordal third (*) differed across stimuli. FFRs (collapsed across groups) did not differ between active vs. passive conditions in the time **(C)** nor frequency **(D)** domains.

**Figure 3.**
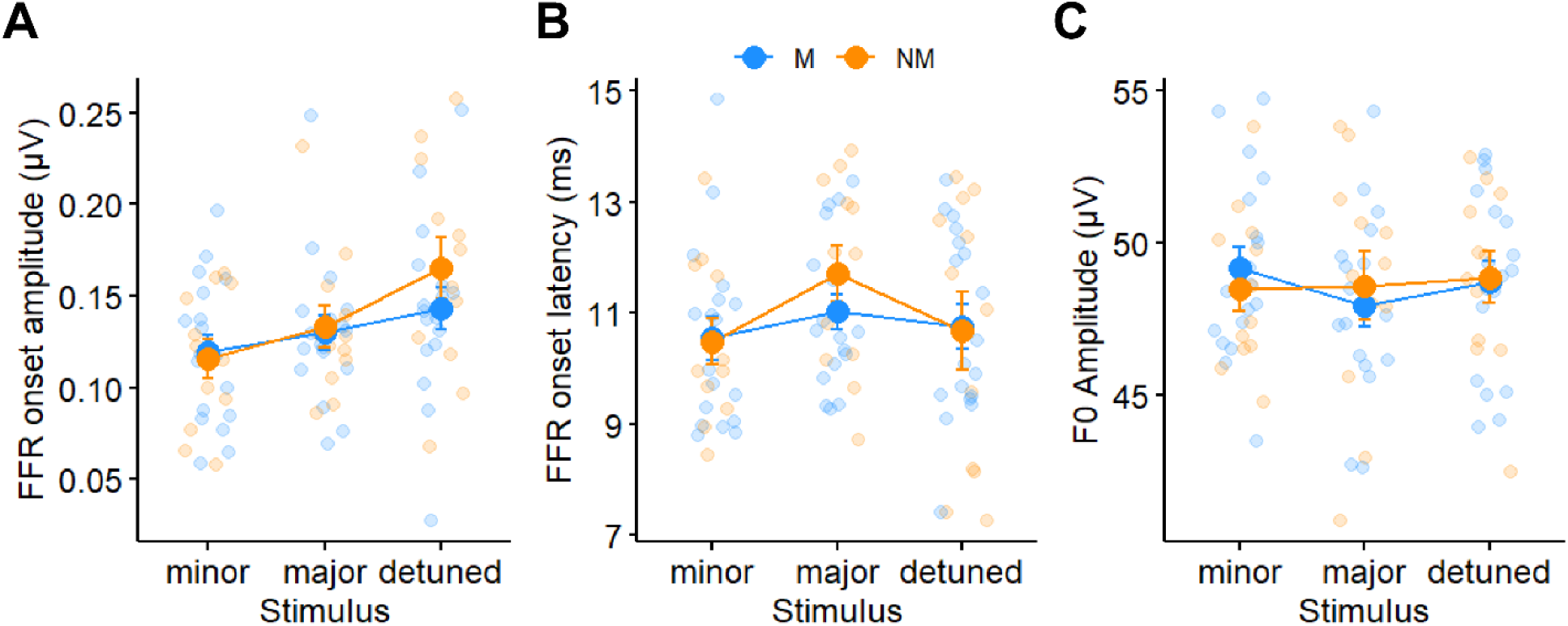
FFR latency and amplitude characteristics. **(A)** FFR onset amplitude varied with chord stimulus but not attention or group. **(B)** FFR onset latency and **(C)** FFR F0 amplitude were invariant both within and between participants.

In the spectral domain, F0 amplitudes to the chordal third were isolated from FFR waveforms (**Figure 2B**). F0 amplitudes did not vary with attention condition (**Fig. 2D**) [*F*(1, 65) = 0.23, *p* = 0.63, 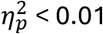], stimulus (**Fig. 3C**) [*F*(2, 65) = 0.52, *p* = 0.60, 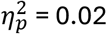], group [*F*(1, 13) = 0.0009, *p* = 0.98, 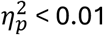], nor their interactions (all *ps* > 0.1).

### Cortical ERP responses

**Figure 4** depicts cortical ERPs for musicians and nonmusicians in both active and passive conditions. An ANOVA on P2 latencies revealed a sole interaction effect between condition and group (**Fig. 5A**) [*F*(1, 65) = 4.20, *p* = 0.045, 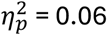]. This was attributed to earlier latencies for the passive relative to active condition in nonmusicians only (**Fig. 4B**; *p* = 0.036). In contrast, musicians showed invariant P2 latencies with attention manipulation (**Fig. 4A**; *p* = 0.54). All other main and interaction effects were non-significant (all *ps* > 0.12).

**Figure 4.**
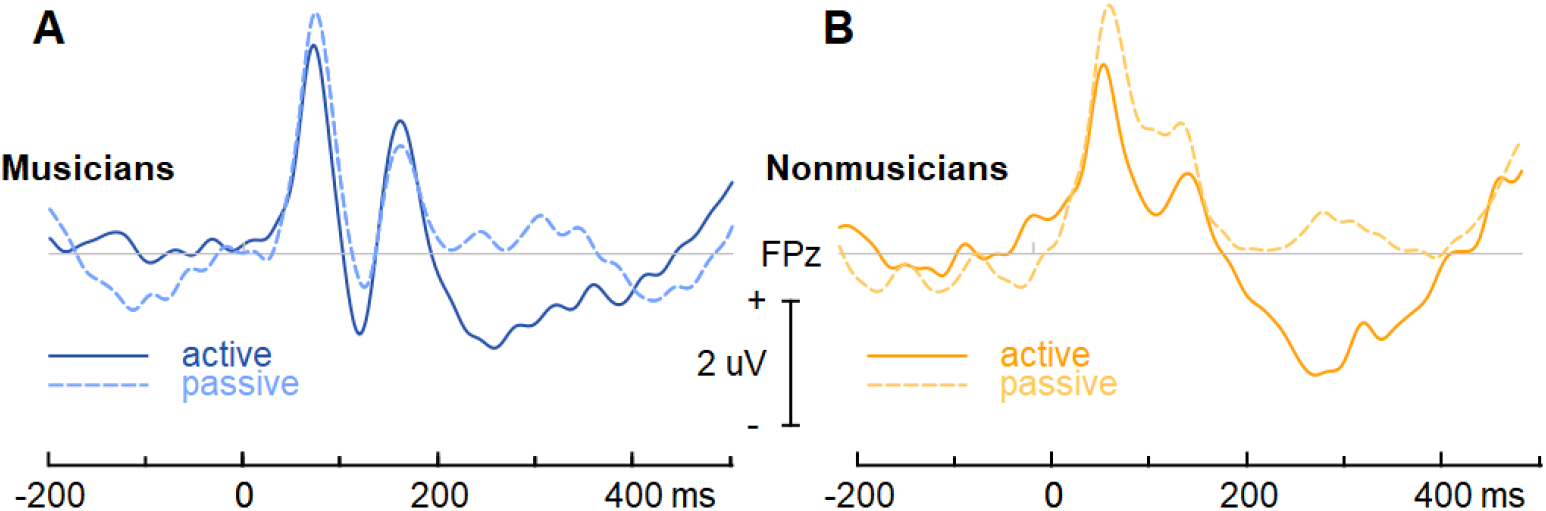
Group averaged ERP waveforms during active and passive conditions. (**A**) Musicians P2 latencies did not differ with attention. (**B**) Nonmusicians showed earlier and stronger P2 responses in the passive compared to active condition.

**Figure 5.**
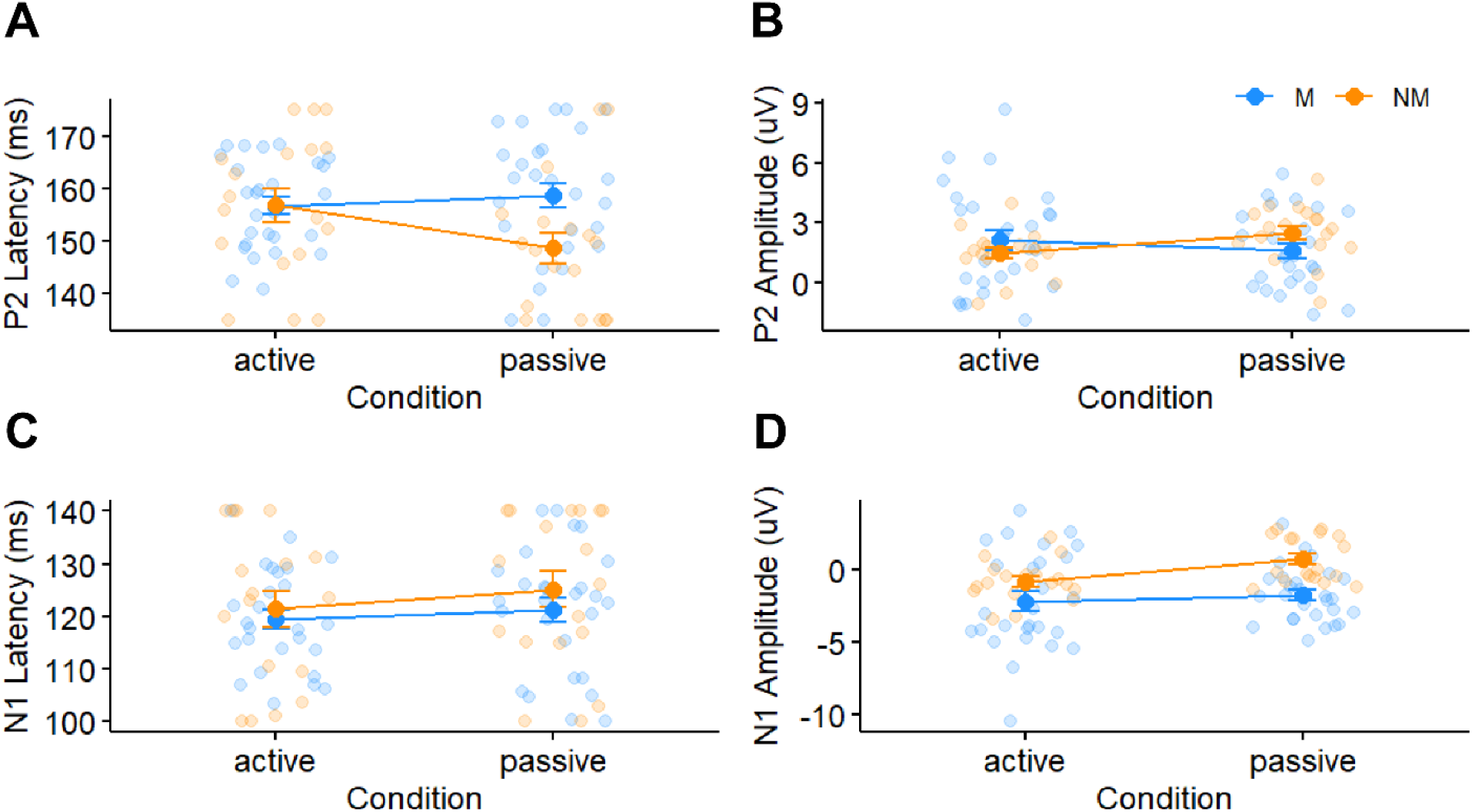
ERP latency and amplitude characteristics. (**A**) P2 latency and (**B**) amplitudes varied with attention and group. **(C)** N1 latency was invariant. **(D)** N1 amplitudes showed a stronger negativity with attention.

The ANOVA on P2 amplitudes revealed an interaction between condition and group (**Fig. 5B**) [*F*(1, 65) = 7.16, *p* = 0.0094, 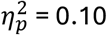]. The interaction was driven by higher amplitudes for the passive relative to active condition in nonmusicians (*p* = 0.029). Paralleling latency, response amplitudes were invariant to attention in musicians (*p* = 0.14). All other main and interaction effects were non-significant (all *ps* > 0.10).

In contrast to P2, N1 latencies were invariant to all experimental manipulations (**Fig. 5C**). However, N1 amplitudes varied with a main effect of attention (**Fig. 5D**) [*F*(1, 65) = 6.07, *p* = 0.016, 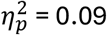]; N1 amplitudes were stronger (i.e., more negative) in the active than passive condition. All other main and interaction effects were non-significant (all *ps* > 0.08).

### Brain-behavior relationships

**Figure 6A** displays the mean Euclidian distances between all pairwise ERPs (N1-P2 amplitudes) for each group. This measure reflects the degree to which the neural responses differentiated musical chords. Musicians demonstrated larger differences in ERP amplitudes across chords than musicians suggesting more distinct neural responses [*t*(13) = 2.59, *p* = 0.022]. **Figure 6B** shows the correlation between neural ERP distance and behavioral information transfer. The positive association between brain and behavioral responses suggests that greater cortical neural differentiation of chordal stimuli (as in musicians) was related to better behavioral IT (r=0.59, *p* = 0.02), and thus less confusability of the chords. In contrast to the cortical ERPs, none of the FFR measures including onset and spectral amplitudes/latencies showed this brain-behavioral correspondence (all *p*s > 0.05; data not shown).

**Figure 6.**
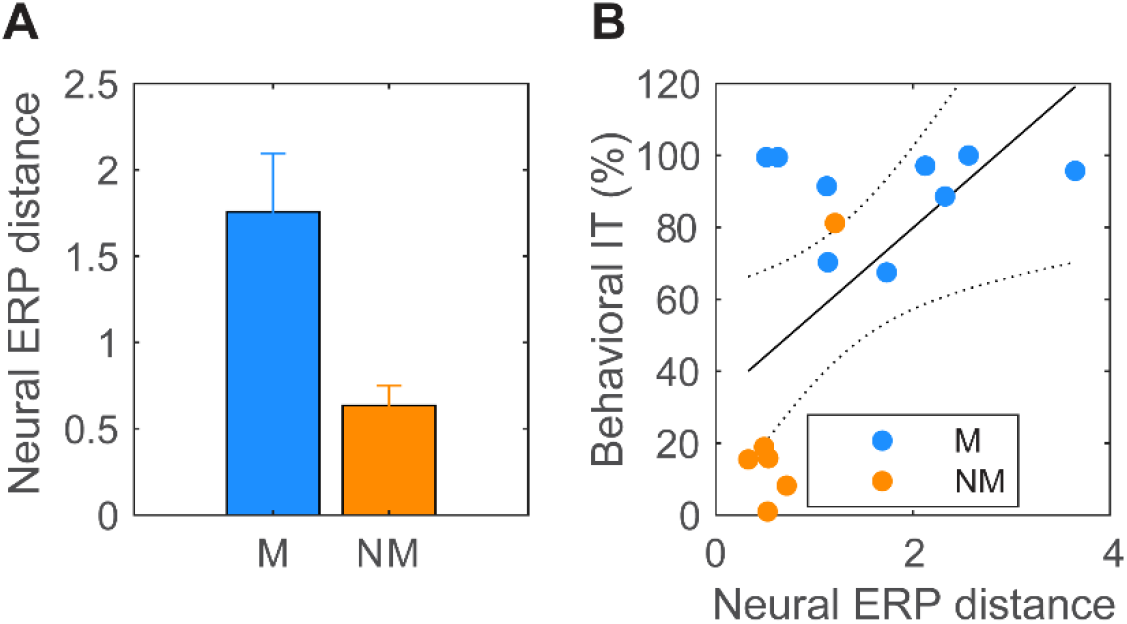
Brain behavior relation in the differentiation of musical chords. **(A)** ERP distance between chords (Euclidian distance between all pairwise N1-P2 amplitudes) is larger in musicians, indicating more salient neural responses across triads. **(B)** ERP distance between chords correlates with behavioral IT indicating a brain-behavior correspondence between perceptual and neural (cortical) differentiation. No such brain-behavior correspondence was observed for brainstem FFRs. Error bars: ± 1 S.E.M.

## DISCUSSION

By recording subcortical FFRs and cortical ERPs to harmonic chordal stimuli during active and passive conditions in musicians and nonmusicians, we found that: (i) subcortical responses did not vary with musicianship or attention; (ii) musicianship and attention strongly modulated cortical responses; and (iii) better cortical differentiation of chord stimuli in musicians corresponded with their better identification (e.g., less confusions) of musical sounds behaviorally.

### Subcortical responses are largely invariant to musicianship and attention

As expected, we found FFRs faithfully encoded stimulus acoustics, representing differences in frequency of the chordal third. Similarly, FFR onset amplitude changed with stimuli, suggesting this feature is sensitive to changes in the chordal third in otherwise identical simultaneous chords.

Prior work using musical stimuli to elicit FFRs has been done during passive listening [1,2,39,56-60]. Yet, attention has been shown to modulate speech-evoked FFRs [22,23,37,61-63]. To determine the role of attention in subcortical music encoding, we recorded FFRs in both active and passive listening conditions. However, we did not observe an overall difference in FFRs with attention (though attention did interact with stimulus as discussed below). The lack of global attention effect could be due in part to our stimulus parameters. FFRs are generated by several cortical and subcortical sources whose engagement varies in a stimulus-dependent manner. Stimuli with high F0s elicit FFRs that are dominated by brainstem contributions [35,64-66]. The high F0 of our stimuli (≥220 Hz), used to intentionally minimize cortical contributions to the FFR, attenuates attentional modulations from the cortex, which may drive previously reported FFR enhancements [67,68]. Though attention may also enhance subcortical FFRs, this modulation is likely dependent on top-down feedback from cortex [37], the strength of which varies between individuals [69]. Consequently, it is also possible that listeners in our sample had weak cortical feedback, subduing any attentional gain to the FFR that is sometimes measurable (Price & Bidelman, 2021).

Interestingly, we did observe an interaction between attention and stimulus conditions on FFR onset amplitude. We found larger differences in onset amplitude with attention for the major vs. minor and detuned chords. This suggests some degree of stimulus-dependent attention effect at the brainstem level. Prior work has demonstrated FFR pitch salience corresponds with perceptual ratings of consonance with major chords having highest pitch salience and consonance ratings [70]. The interaction we observed could reflect stimulus-dependent attentional enhancements corresponding to consonance or some salient feature of major, but not minor or detuned, chords (e.g., perceptual brightness).

Surprisingly, we did not observe changes in FFRs with musicianship. Several studies have reported enhanced FFRs to both speech [71-75] and musical (usually isolated notes) stimuli [1,5,57] in musicians. However, many of these studies used stimuli with low F0, potentially allowing for greater cortical contributions to the FFR. Indeed, musician’s enhanced FFR might be attributable to larger cortical rather than subcortical phase-locking (Coffey et al, 2016). Our use of higher F0 stimuli here intentionally elicits FFRs dominated by brainstem generators, which would render any cortical effects of long-term musical training unobservable. The findings here echo those of MacLean et al. [21] who also reported similar FFRs between musicians and nonmusicians for high-frequency speech stimuli. Our results support the notion that FFR enhancements traditionally reported for musicians might be influenced more by cortical rather than subcortical neuroplasticity.

### Cortical responses vary with attention and musicianship

In stark contrast to brainstem FFRs, attentional effects were much stronger in the cortical ERPs. Consistent with prior literature on ERP attentional effects, we observed large enhancement of N1 amplitudes in the active compared to passive condition [24,44,45,76]. Comparisons between groups showed earlier and stronger P2 responses in nonmusicians during passive listening, whereas musicians’ P2 was invariant to attentional manipulation. P2 is a wave that occurs relatively early after stimulus onset (∼150 ms) and is often associated with perceptual auditory object formation [77]. Musicians’ similar P2 profiles in passive and active conditions suggest automaticity in early sound processing. Contrastively, attention seemed to have a stronger effect on sensory coding in nonmusicians. Though we may have expected earlier and stronger P2 during active listening [78], nonmusicians’ P2 was reduced and later in the active compared to passive condition. Nonmusicians’ P2 reductions in the active condition could result from habituation with short-term learning of the musical sounds. This parallels similar habituation effects observed during speech-learning [21,79,80].

### Cortical stimulus differentiation mirrored behavioral discrimination

We observed a relationship between neural stimulus differentiation (assessed through pairwise N1-P2 amplitude differences) and behavioral information transfer. In other words, the degree to which listeners *behaviorally* distinguished triads was directly related to their *neural specificity* to stimuli primarily at a cortical (rather than subcortical) level. These results suggest that cortical levels of processing more closely support the behavioral distinction of chordal stimuli. Moreover, the higher predictive power of ERPs vs. FFRs we find in predicting listeners’ behavioral chord confusions implies that cortical neural representations convey a closer readout to the ultimate percept.

Our cortical findings support previous ERP studies examining stimulus differentiation in musicians vs. nonmusicians using the mismatch negativity (MMN), a pre-attentive potential indexing the brain’s automatic change detection for sounds [3,30,81,82]. Musicians’ enhanced behavioral pitch discrimination abilities are mirrored in their enhanced MMNs to speech pitch and timbre [83], detuned chords [30], and changes in the chordal third of musical triads [3]. These findings suggest long-term music training with complex pitched stimuli facilitates the development of superior auditory discrimination at both the neural and behavioral levels [82]. Here, we demonstrate musicians’ enhanced behavioral identification of musical chords corresponded with greater cortical discrimination in the ∼150 ms time window. This provides further evidence that music training strengthens cortical specificity for musically relevant sounds.

## CONCLUSION

Our results suggest that long-term music training strengthens the brain’s encoding of musical sounds, particularly at a cortical, rather than subcortical level of processing. We also find more attention-related changes in the ERPs of nonmusicians vs. musicians. We propose that these findings add to existing evidence for more automated neural processing of musical chords at higher levels which do not depend on attention. Additionally, our findings demonstrate that *cortical* specificity to musical triads is better in musicians and predicts their enhanced *behavioral* identification. This argues for a close relationship between high-level neural and behavioral distinction of musical sounds.

